# The context-dependent nature of the neural implementation of intentions

**DOI:** 10.1101/401174

**Authors:** Sebo Uithol, Kai Görgen, Doris Pischedda, Ivan Toni, John-Dylan Haynes

## Abstract

Many studies have identified networks in parietal and prefrontal cortex that are involved in intentional action. Yet, knowledge about what these networks exactly encoded is still scarce. In this study we look into the content of those processes. We ask whether the neural representations of intentions are context- and reason-invariant, or whether these processes depend on the context we are in, and the reasons we have for choosing an action. We use a combination of functional magnetic resonance imaging and multivariate decoding to directly assess the context- and reason-dependency of the processes underlying intentional action. We were able to decode action decisions in the same context and for the same reasons from the fMRI data, in line with previous decoding studies. Furthermore, we could decode action decisions across different reasons for choosing an action. Importantly, though, decoding decisions across different contexts was at chance level. These results suggest that for voluntary action, there is considerable context-dependency in intention representations. This suggests that established invariance in neural processes may not reflect an essential feature of a certain process, but that this stable character could be dependent on invariance in the experimental setup, in line with predictions from situated cognition theory.

## 1. Introduction

Intentions are believed to operate at the interface of thought and action (Bratman, 1987), and therefore have to translate cognitive states into detailed motor programs (Mylopoulos & Pacherie, 2017). They are assumed to be the end state of a decision process, and to be the primary cause of the subsequent action. Due to this assumed central role intentions play in cognition, they have a rich history of scientific investigation, going back to William James (1890), and Kornhuber & Deecke (1965).

Even though the (folk) psychological notion of “intention” assumes a homogeneous or unitary process, neuroscientific evidence suggests that intentional actions consist of multiple components and that various brain structures is involved. For example, human stimulation studies show that the urge to move can be contrasted with the desire to move (Desmurget & Sirigu, 2012). The former is evoked when the medial prefrontal cortex—specifically the supplementary motor area (Fried et al., 1991)—is stimulated, whereas a desire to make a movement (in less detail) is evoked upon stimulation of the inferior parietal lobule (Desmurget et al., 2009).

In order to accommodate the multitude of brain areas involved in voluntary action, and the various roles intentions are thought to play, Jahanshahi (1998) suggested that intentions consist of multiple components, including a “what to do” component, a decision when to act, and an inhibitory processes. A similar decomposition can be found in Brass and Haggard’s (2008) “what”, “when”, and “whether” model of intentional action.

These findings and models explicate the stages and structures involved in intentional action. Yet, what exactly is encoded in these neural representations of intentions has received less attention. The standard, but mostly tacit assumption is that the content of intentions are representations of a desired outcome (Haggard, 2005; Uithol, van Rooij, Bekkering, & Haselager, 2012). Alternatively, embodied approaches to cognition suggest another possibility. In real life, our action decisions are often strongly connected to the context in which they are made, that is the immediate environment of the intended action. The context may actively suggest or afford certain types of behavior (Gibson, 1979). Consequently, part of what is needed to complete an action may not be represented in a discrete and invariant intention, but be guided by contextual input from the environment (Clark, 1997). Along these lines, recent findings suggest that action decisions and action planning processes cannot be separated (Andersen & Cui, 2009; Cisek, 2007; but see Bennur & Gold, 2011). So, when movement-planning is necessarily context-dependent (otherwise it would not lead to successful behavior), and when action-selection and movement planning are tightly interwoven, then action decision processes may be much more context-dependent than commonly assumed.

Down similar lines, different reasons for performing an action have been shown to result in different kinematics (to the extent that they can be picked up by a human observer: Ansuini, Cavallo, Bertone, & Becchio, 2014), suggesting that different reasons may be responsible for differences in the processes underlying intentional actions.

Here we directly test the reason- and context-dependency of action decisions. Specifically, we investigate whether the same action decision made for different reasons or in different contexts is accompanied by invariant neural patterns (as indexed by multivariate analyses of fMRI data). Embodied approaches would predict that the same action decision in different contexts might exhibit radically different patterns. More traditional approaches would allow for some contextual variation, but would still predict a ‘core representation’ with the chosen action as its content. Both embodied and traditional approaches would predict that changing the reasons for choosing an action has notable (see above), but marginal effects on the representation of action intentions.

In this study participants formed action intentions based on specific reasons and in specific contexts. Given a certain reason only one of the actions was reasonable. However, participants were not explicitly instructed by us to make a particular choice. We chose to use such semi-free decisions for two reasons: First, in real life we virtually never form intentions that are completely free from reasons to act. In general we form intentions because we want to achieve a certain goal, and that goal is often—if not always—embedded in a context^1^. Second, this setup allows us to check that participants perform the task (by assessing their response) and to balance the number of trials per condition.

## 2. Methods

### 2.1 Participants

Thirty participants took part in this study (mean age 26.1 years, 21 female). All participants had normal or corrected-to-normal vision and were right-handed according to the Edinburgh handedness assessment (Oldfield, 1971). Participants had no history of neurological or psychiatric disorders, and gave written informed consent. All participants mastered the German language at a native level, and received 20€ for participation. The study was approved by the local ethics committee.

### 2.2 Experimental setup

Stimuli were presented using PsychoPy version 1.83.03 (Peirce, 2007). They were projected onto a screen at the back of the scanner that was visible through a mirror mounted to the MR head coil. Each trial (Fig. 1) started with an image depicting a contextual setting (either a breakfast or a supermarket context), presented at the center of the screen (factor “context”). Beneath the picture a one-sentence explanation of a situation was presented in German (factor “reason”). For example, “You have poured milk in your glass and you’ve put the lid back on”, which suggests placing the box (action) in order to do away with it (reason) in a breakfast context. This combination of picture and sentence was presented for 4000 ms followed by a 6000 ms decision delay with an empty gray screen. After that, the response screen was presented, which always contained the two options “open” and “place”. On half of the trials these options were presented as words, on the other half of the trials as pictograms, to prevent participants from anticipating specific visual input. The two possible answers were each at different sides of the screen, and participants indicated the correct answer by pressing a button with either their left or right index finger. The side at which each answer appeared was randomized to prevent motor preparation. Participants were asked to respond within 2000 ms after which responses were considered invalid. The response was followed by a 2000 ms intertrial interval in which participants could prepare for the next trial.

**Figure 1.**
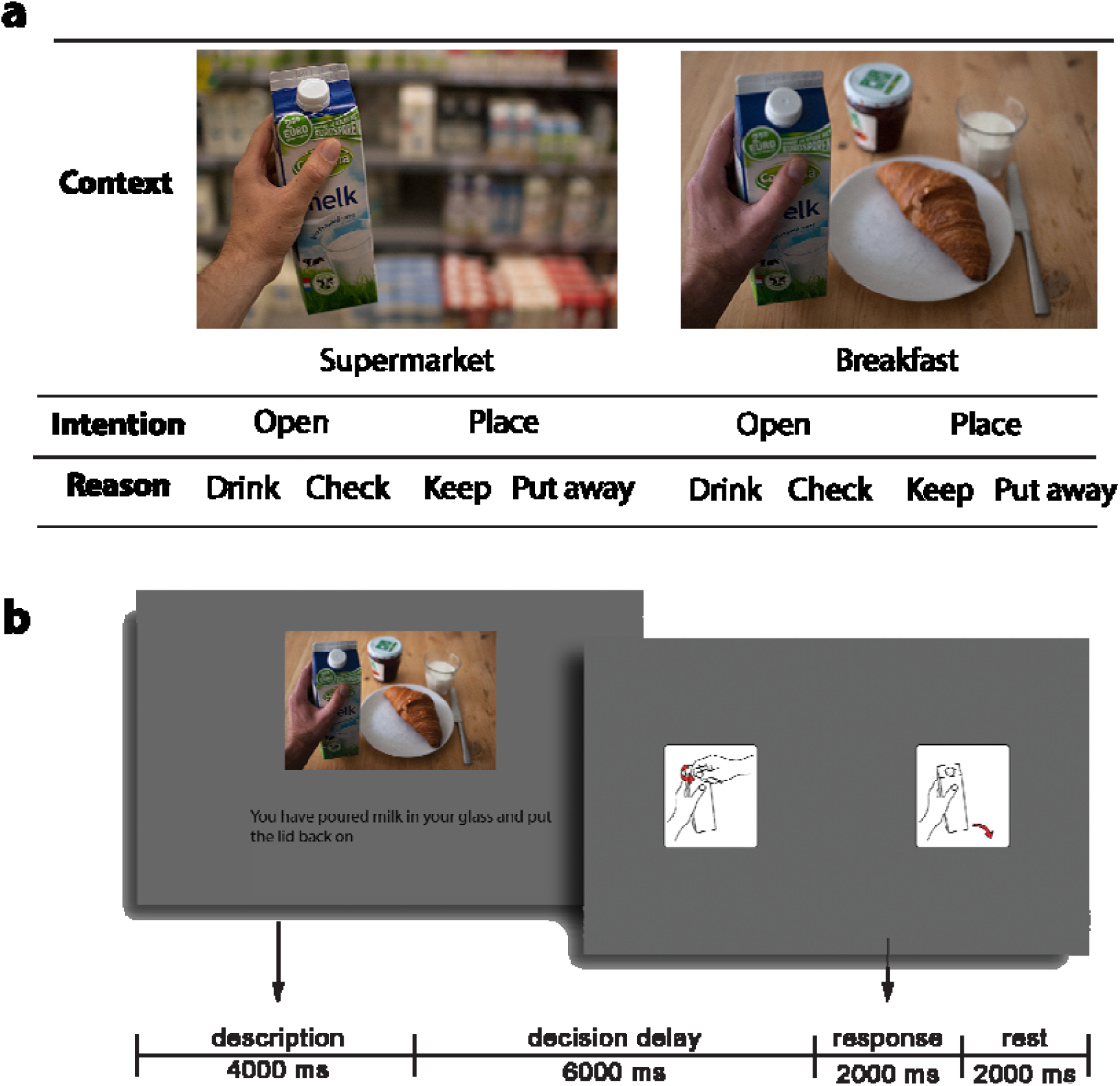
Experimental design. Panel **a:** schematic overview of the eight experimental conditions. The photos here are examples; various photos were used. Panel **b**: Graphical representation of one trial. The originally German sentence has been translated to English. During the delay and the intertrial interval an empty gray screen was presented.

There were two contexts (supermarket vs. breakfast), two action options (open vs. place) and two reasons for choosing one action or the other, resulting in a 2 x 2 x 2 design. As the reasons for opening were different from the reasons for placing, this factor was nested. Each of the eight conditions was repeated four times within one run. Participants performed five runs (separated by short breaks), thus each condition was repeated 20 times, resulting in a total of 160 trials. Participants received instructions and a short training session prior to the experiment. The total duration of the experiment, without setting up and training, was approximately 40 minutes.

Trial order was randomized, and the experimental design was assessed using the Same Analysis Approach (Görgen, Hebart, Allefeld, & Haynes, 2017), to test for unintended regularities in the design that could bias the machine-learning classifier (see below). No such regularities were found.

### 2.3 Image acquisition

A 3-Tesla Siemens Trio (Erlangen, Germany) scanner with a 12-channel head coil was used to collect functional magnetic resonance imaging data. In each run, 266 T2*-weighted echo-planar images (EPI) were acquired (TR = 2000 ms, TE = 30 ms, flip angle 90°). Each volume consisted of 33 slices, separated by a gap of 0.75 mm. Matrix size was 64 × 64, and field of view (FOV) was 192 mm, which resulted in a voxel size of 3 × 3 × 3.75 mm. The first three images of each run were discarded. Additionally, a structural, T1-weighted image was collected for anatomical localization.

### 2.4 Data analysis

The EPI images were preprocessed using SPM12 (http://www.fil.ion.ucl.ac.uk/spm). The images were realigned, unwarped and slice-time corrected. Next, we estimated a general linear model (GLM; Friston et al., 2004) with 12 regressors corresponding to the 8 conditions in the design plus the stimulus pictures (in order to minimize a possible effect of visual information on the performance of the classifier in the subsequent analyses). Regressors were modeled as a box-car, encompassing the delay period (4-10 s from trial onset, see Figure 1) and convolved with a canonical hemodynamic response function. We also included 6 movement parameters as regressors of no interest. The condition-, voxel-, and run-wise parameter estimates of the resulting GLM were subsequently used as input for the multivariate analyses.

#### 2.4.1 Multivariate decoding

We employed a multivariate pattern analysis (MVPA) using The Decoding Toolbox (TDT; Hebart, Görgen, & Haynes, 2015). A searchlight classifier using libSVM (Chang & Lin, 2011) was trained to classify the action (open vs. place) for one specific reason-context combination based on data from four of the five runs. The searchlights had a radius of 12 mm and were restricted within a whole-brain mask. This classifier was then subsequently tested on all four reason-context combinations in the remaining fifth run, constituting a 5-fold run-wise cross-validation. The four test combinations were (1) classifier trained and tested on the same reason and the same context (‘SameReasonSameContext’), (2) classifier trained and tested on the same context but on a different reason (‘CrossReason’), classifier trained and tested on the same reason but in a different context (‘CrossContext’), and (4) classifier trained and tested on different reasons and different contexts, (‘CrossReasonCrossContext’, see Fig. 2). This procedure was repeated for each of the four context-reason pairs and using data from each of the five runs as test set once. The classification accuracies from these four context-reason pairs and the five runs were averaged.

**Figure 2.**
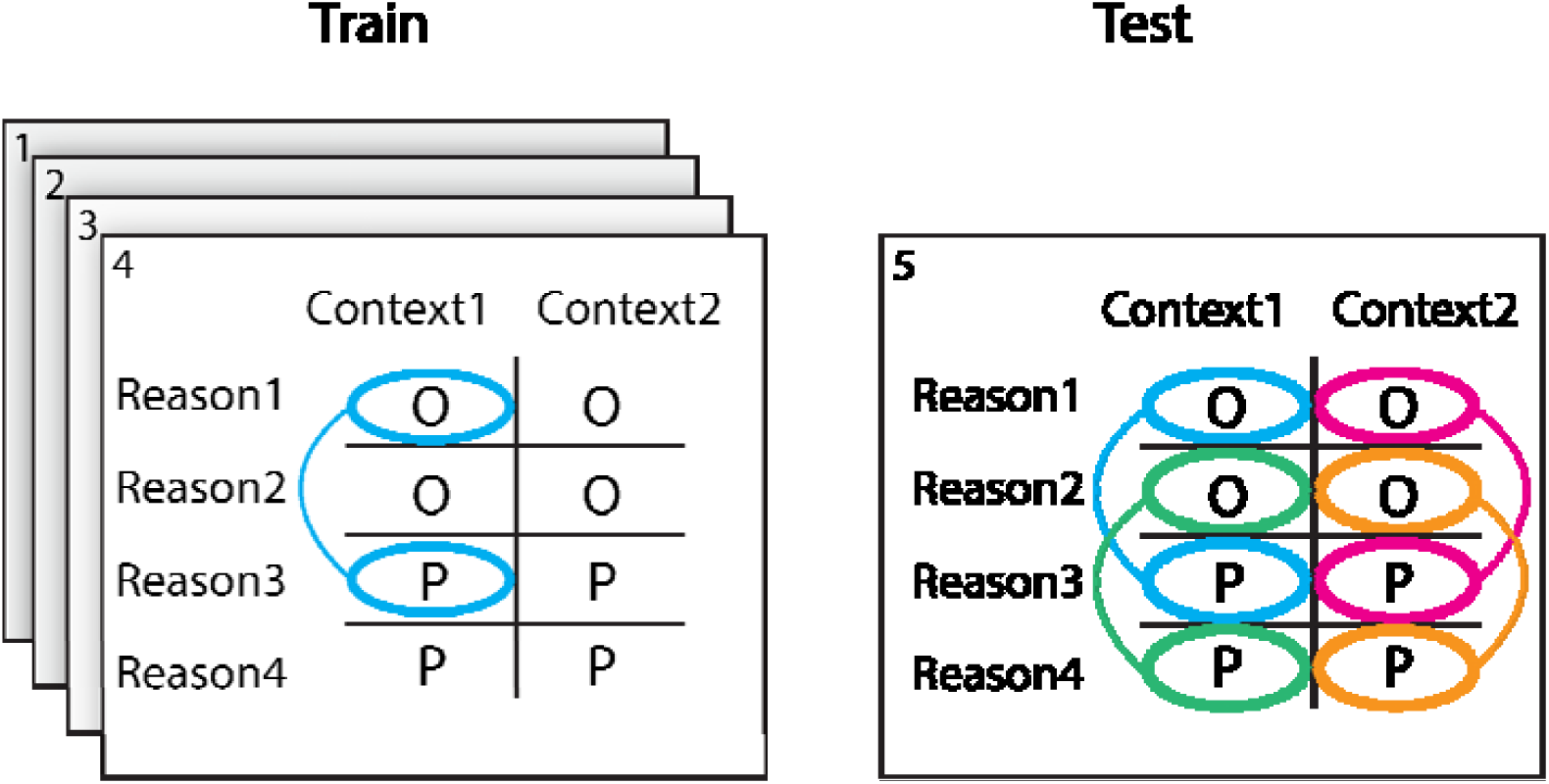
The training and testing procedure. For training and testing we employed a leave-one-run-out procedure. A classifier was trained on four of the five runs to distinguish between the intention to open and to place, and subsequently tested on the left-out run in four different settings: 1) SameReasonSameContext (blue), 2) CrossReason (green), 3) CrossContext (magenta), and 4) CrossReasonCrossContext (orange).

In order to increase sensitivity we restricted our crucial generalization analyses to regions that encoded intention-related information for the same context and the same reason. Thus, we created region of interests (ROIs) based on successful SameReasonSameContext classification (combination 1 in Fig. 2). These ROIs were normalized to MNI-space (3^rd^ degree B-Spline interpolation) using the structural images, and smoothed with a 2×2×2mm FWHM kernel. A second level analysis over subjects was used to find cortical regions that allowed for above-chance decoding of action intentions. To avoid circularity for the estimate of the SameReasonSameContext condition (Kriegeskorte, Simmons, Bellgowan, & Baker, 2009; Vul, Harris, Winkielman, & Pashler, 2009) the ROIs for each participant were created using a leave-one-participant-out protocol (e.g., the ROIs of participant 1 were created using the decoding results of all subjects but participant 1). For this, a threshold of p<0.001, family-wise error corrected at cluster level, was used (Flandin & Friston, 2017). The ROIs covered a substantial part of the cortex, including visual, parietal, frontal, prefrontal and temporal regions (see Fig. 3, left panel). As we were not interested in visual decoding (which could be driven by regularities in the stimuli), only parietal, premotor, prefrontal, and cerebellar areas were used for the following analyses. The continuous ROI was split into four functional anatomical ROIs each subject (see Fig. 3, right panel; the exact boundaries of these ROIs varied per participant). All ROIs were present on both hemispheres, but typically larger on the left one.

**Figure 3.**
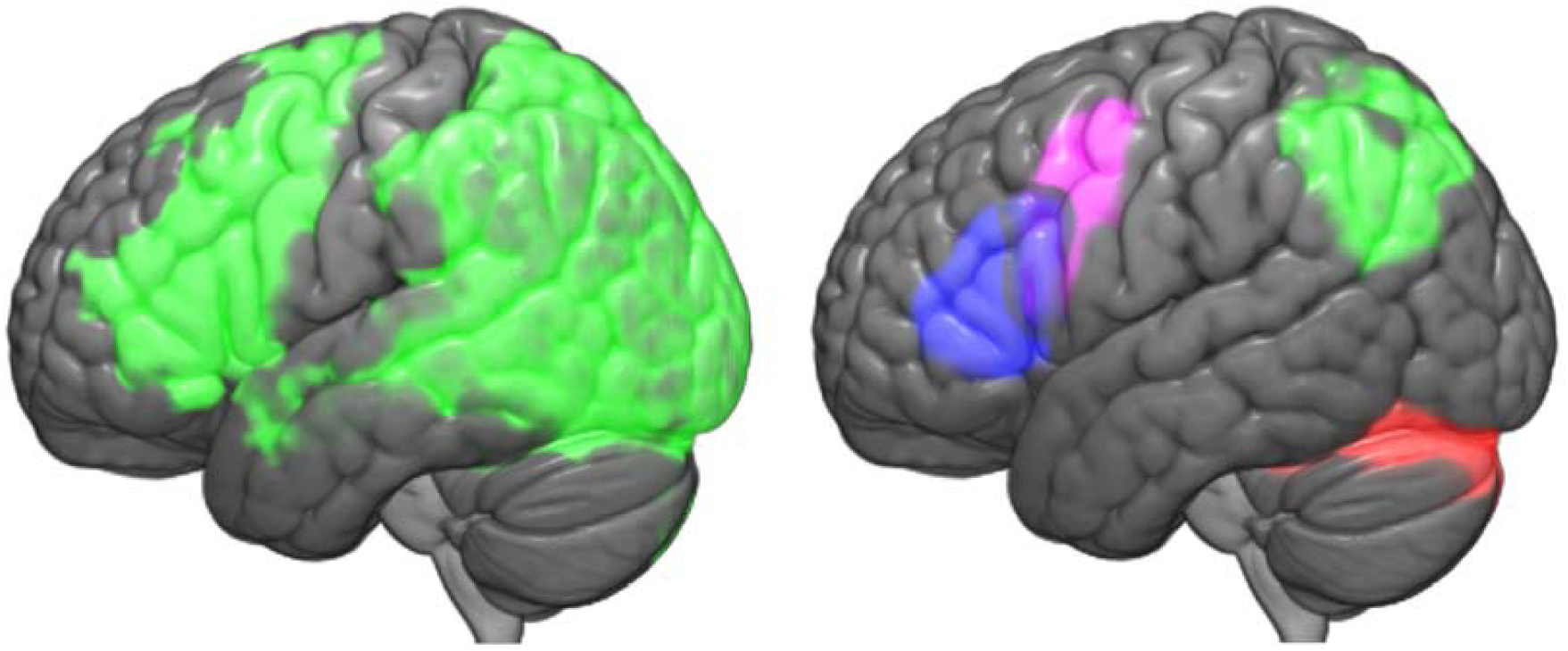
**Left picture** shows the (smoothed) regions that allowed for above-chance decoding in the SameReasonSameContext condition based on a second-level analysis including all subjects. **Right picture** shows the four ROIs used for subject 1 during the ROI analysis: prefrontal (blue); premotor (purple); parietal (green); and cerebellum (red). To avoid circularity in the SameReasonSameContext, ROIs were created for each subject from the classification results of all other subjects (here: all except subject 1).

**Figure 4.**
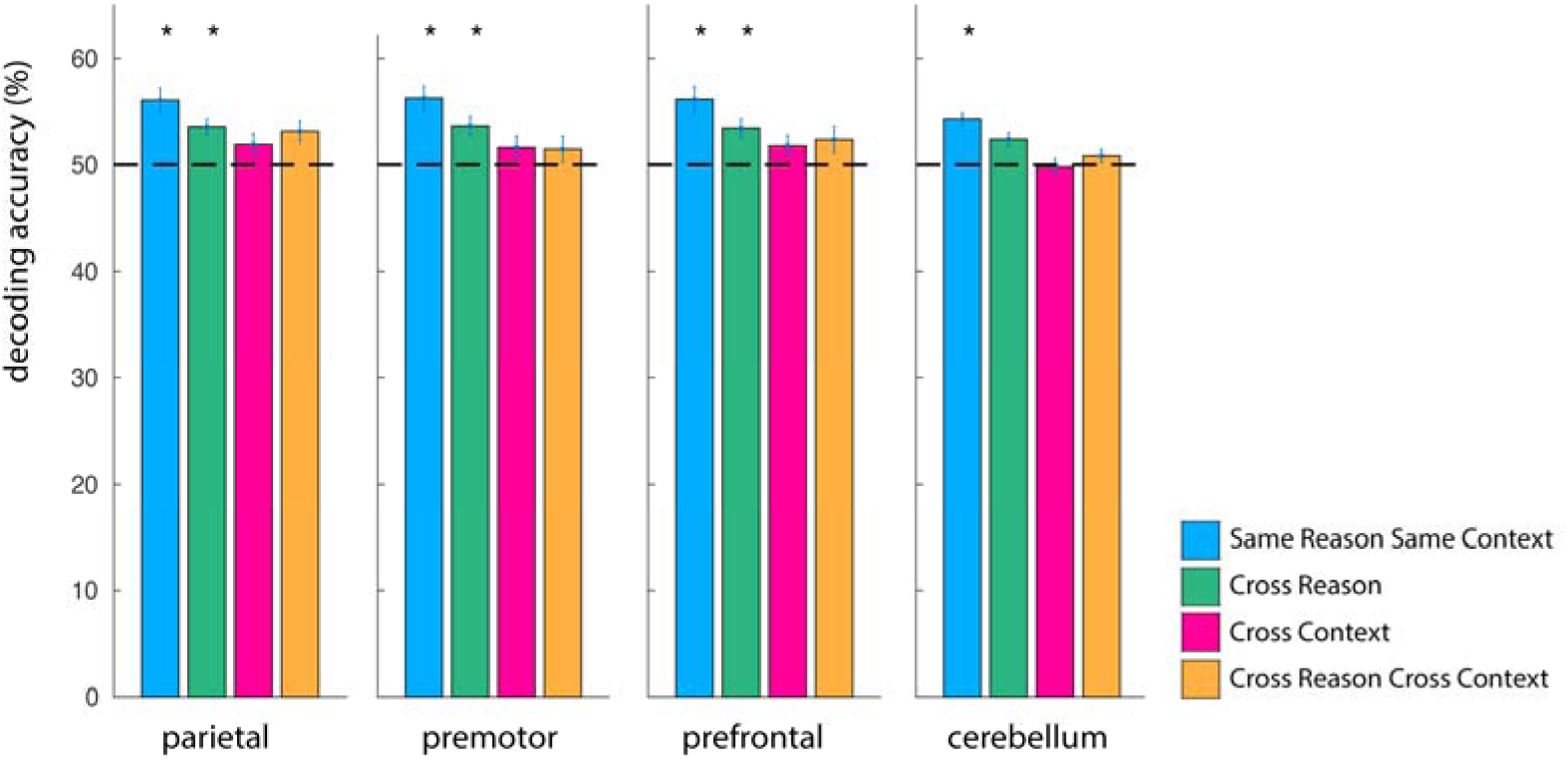
Results of the decoding per ROI. Asterisks denote values being significantly (p<0.05, Bonferroni-corrected for multiple comparisons) above chance level (50%, horizontal dashed line). Error bars represent standard error of the mean.

These ROIs were used to compare the four different (cross-)classification performances. To this end we used a whole-brain searchlight approach on unsmoothed images in native space (again using a searchlight with a 12mm radius). We generated individual ROIs using the inverse transformation matrix that was generated during normalization. Subsequently, the classification accuracies were extracted from the whole-brain searchlight analysis for each of the ROIs. The group average classification in the four conditions (SameReasonSameContext, DifferentReason, DifferentContext and DifferentReasonDifferentContext) was compared in each of the four ROIs.

## 3 Results

### 3.1 Behavioral results

Trials that contained responses that were either incorrect or that occurred outside of the instructed response window of 2000ms were discarded. Only participants with two or more (out of four) correct trials on each of the eight conditions in each run were included in the analyses reported below. Four participants were excluded based on this criterion. The remainder of the subjects gave on average the matching answer on 94% of the trials (standard deviation 6.5). Since there was a 6000ms delay between stimulus presentation and answering screen, we assume that no meaningful information can be drawn from the reaction times. This is confirmed by the repeated-measure ANOVA, showed no effect of condition on reaction time (p > 0.6).

### 3.2 Classification results

The SameReasonSameContext condition showed significant information in all ROIs (classification accuracies: parietal: 56.0%; premotor: 56.2%; prefrontal: 56.2%; cerebellum: 54.3%; p<0.001, significance threshold after correction for multiple comparisons (four conditions, four ROIs), results in α= 0.003). Please note that while the SameReasonSameContext condition was used to define the ROIs, the ROI definition for each participant was performed based on data from all other subjects and thus voxel selection and test-set classifier performance were independent. Furthermore, within a participant the classification was performed in a cross-validated fashion with a left-out run as test data set. CrossReason decoding accuracy was smaller than SameReasonSameContext accuracy, but still significantly above chance level (parietal: 53.6%; premotor: 53.6%; prefrontal: 53.5%; cerebellum: 52.4%, p<0.001 for all ROIs except the cerebellum: p < 0.005, α=0.003). CrossContext decoding was not significant in any of the four ROIs (parietal: 51.9%; premotor: 51.6%; prefrontal: 51.8%; cerebellum: 49.8%, p > 0.05 for all ROIs, α= 0.003), neither was CrossReasonCrossContext decoding (parietal: 53.2%; premotor: 51.4% prefrontal: 52.3%; cerebellum 50.1%, p > 0.003; 0.1; 0.05; 0.1 respectively, α= 0.003).

We also assessed whether the lack of significant decoding in CrossContext conditions was due to context-sensitivity of neural information or, alternatively, to the lack of statistical power. This was assessed using Bayesian statistics, as implemented in JASP (JASP Team, 2017). A Bayes factor BF_10_ was calculated using a Bayesian repeated-measures ANOVA over classification accuracies. The BF_10_ informs how more likely Hypothesis 1 (there being an effect) is than Hypothesis 0 (there is no effect). Following Lee & Wagenmakers (2014), we consider a log(BF) between 0 and 1 to be anecdotal; between 1 and 2 to be moderate; between 2 and 3.5 strong, between 3.5 and 4.5 to be very strong, and larger than 4,5 as extreme. This analysis shows a moderate to extreme effect for context, but not for reason for all ROIs, (see Table 1 for the (natural) logarithm of the Bayes Factors).

**Table 1.** The natural logarithm of the Bayes Factors (BF10) of the Bayesian repeated measures ANOVA for information on the action intentions vs. no information in the different ROIs. Negative values (in red) indicate evidence for the null hypothesis.

| | Log($BF_{10}$) ANOVA Context | Log($BF_{10}$) ANOVA Reason |
| --- | --- | --- |
| **Parietal** | 1.9 | -1.4 |
| **Premotor** | 2.9 | -0.9 |
| **Prefrontal** | 1.0 | -1.1 |
| **Cerebellum** | 7.3 | -1.4 |

Based on the significant context effect of the ANOVA we performed separate t-tests for each individual condition. Table 2 shows the logarithms of the Bayes Factors for the different conditions in the different ROIs. Negative values (printed in red) indicate evidence for the null hypothesis (no classificatory information is available in the ROI).

**Table 2.** Logarithms of the Bayes Factors (BF10) for information on the action intentions vs. no information in the different ROIs. Negative values signify evidence for the null hypothesis (i.e., no information is present in the neural activity pattern).

|  | Log(BF<sub>10</sub>) T-Test<br>SameReasonSame<br>Context | Log(BF<sub>10</sub>) T-Test<br>CrossReason | Log(BF<sub>10</sub>) T-Test<br>CrossContext | Log(BF<sub>10</sub>) T-Test<br>CrossReasonCross<br>Context |
| --- | --- | --- | --- | --- |
| **Parietal** | 5.8 | 5.2 | -0.3 | 1.8 |
| **Premotor** | 6.4 | 3.4 | -0.8 | -1.1 |
| **Prefrontal** | 5.6 | 3.2 | -0.4 | -0.8 |
| **Cerebellum** | 10.6 | 2.8 | -0.3 | -0.9 |

It is puzzling why CrossReasonCrossContext in the parietal ROI yields a positive coding result in the Bayesian analysis, whereas CrossContext decoding does not. This suggests that this part of the parietal cortex region contains no information about the action intention when the context is varied, but that it does contain information when the reason is varied in addition. It may be possible that the significant result in the CrossReasonCrossContext condition reflects a false positive (despite the use of Bonferroni correction across tests).

## Discussion

We directly assessed the impact of changing context and changing reasons on the processes underlying action decisions, by means of comparing the decoding accuracy between a within-context and reason to cross-context and cross-reason conditions. When context changed between training and testing the classifier, the decoding accuracy drop to chance level. Changing the reasons for forming a certain intention did not have this effect. This suggests that context plays a crucial role in the processes underlying action intentions, in line with embodied approached to cognition, that emphasize the ‘situatedness’ of cognition (Wilson, 2002).

The notion of a context-invariant encoding of task information in a single brain region has long been challenged: It is known that different variables of tasks might be encoded in different regions across the cortex, supporting a distributed model of task encoding (Jahanshahi, 1998), which is in line with the Multiple Demand Network framework (Fedorenko, Duncan, & Kanwisher, 2013). Furthermore, when preparing actions the information about ‘what’ will be performed and ‘when’ it will be performed is encoded in dissociable brain regions (Soon, Brass, Heinze, & Haynes, 2008; Zapparoli et al., 2018), in line with Brass and Haggard’s (2008) distinction of the ‘what’, ‘when’ and ‘whether’ subprocesses of intentions (see also Krieghoff, Brass, Prinz, & Waszak, 2009; Momennejad & Haynes, 2012). Our results can be interpreted as an extension of this deviation from a singular process, or locus, by showing that the ‘what’ is not a discrete process either, but a process that depends on the context in which the action decision is made.

Furthermore, it has been previously shown that within a single brain region information may be strongly task-dependent (Woolgar, Thompson, Bor, & Duncan, 2011; but see Cole, Ito, & Braver, 2016) and that task information also differs across different sequential stages of task processing (Sigala, Kusunoki, Nimmo-Smith, Gaffan, & Duncan, 2008). Our results extend this previous research by showing that task representations are relatively invariant to certain changes, that is variation in reason, but not to others, that is changes in context. It is known that context affects action responses (Wokke, Knot, Fouad, & Ridderinkhof, 2016), and most scholars would assume that action intentions have context-dependent elements, as otherwise they would not be able to consider contextual factors in establishing appropriate actions. Yet, our results provide evidence for stronger context-dependency. Decoding accuracy was not only significantly lower when testing across context (compared to within context) it was also no longer significantly different from chance level. There are two possible explanations for this result. First, there is nevertheless a common representational core underlying intentions for the same action across contexts, but this core is not accessible via the coarse sampling of neural signal with fMRI (i.e. a false negative). A null-hypothesis is hard to prove, and the absence of proof should not be considered proof of absence. Second, it could mean that there is indeed no invariant representational core at all. The Bayesian statistical analysis provided anecdotal to moderately strong evidence in favor of this hypothesis. In that case action-control processes may perhaps be better understood as a dynamic integration of sensorimotor processes (Gilbert & Fung, 2018; Schurger & Uithol, 2015; Uithol, Burnston, & Haselager, 2014), rather than the formation of an invariant intention that is subsequently translated into the current context (Grafton & Hamilton, 2007; Hamilton & Grafton, 2007; Pacherie, 2008).

Even though we made an effort to make our study more ecologically valid by varying contexts and reasons, we were forced to accept less ecological aspects in our setup: 1) the participants were not really engaged in the context, they had to imagine it and we presented the context by means of a photograph on the screen; and 2) participants did not execute the exact action they had chosen, but they indicated their choice by a button-press. Also, there was a six seconds delay between the onset of the intention formation and the moment the participant could indicate the chosen option. This delay was necessary to ensure a sufficient SNR. We do not have a clear experimental handle on what participants exactly did during this delay, apart from maintaining the chosen option in mind (as reflected in the high accuracy rate in the responses). Participants were asked to imagine themselves in the indicated context, and to imagine the chosen action. In line with this, changing the context had a significant influence on the brain activity during this time frame.

It is an open question to what extent our results, stemming from a task in which meaningful actions in specific contexts were selected, but not executed, can be compared to previous studies in which mostly purposeless actions in a context-free environment were both selected and executed (Gallivan, Johnsrude, & Randall Flanagan, 2015; Gallivan, McLean, Smith, & Culham, 2011a; Gallivan, McLean, Valyear, Pettypiece, & Culham, 2011b). Both approaches are a simplification, but in a different way: Our study is a simplification in the sense that the chosen action is not executed but only imagined, whereas previous studies are a simplification in the sense that the executed actions are lacking an ecological purpose (i.e., an ecologically valid reason) and a meaningful context. Nevertheless, there is some overlap in the cortical areas from which information could be decoded, including ventral premotor, lateral prefrontal and parietal areas. On the other hand, many of the regions that have been previously reported for being involved in action selection are absent in our results: frontal eye fields, dorsal premotor and of course primary motor areas, frontopolar and medial prefrontal cortex, as well as pre-SMA (Gallivan et al., 2011b; Goldberg, 1985; Haynes, Sakai, Rees, Frith, & Passingham, 2007; Momennejad & Haynes, 2012; Soon et al., 2008; Woolgar et al., 2011). These regions are often considered the locus of intentional action (Haggard, 2008). Yet, we found no information in these areas in our study. This discrepancy could be explained in two ways: 1) these areas are actually involved but the activity pattern is not (significantly) different between the two intentions, or 2) since our study did not involve executing the selected action (the button press only related to the chosen action in an arbitrary way) no action-planning component (specific to the chosen action) is involved, which may be necessary for these additional areas to be recruited. On the other hand, a previous study that dissociated decision processes from overt movement (Haynes et al., 2007) did find information in medial PFC. This discrepancy could suggest that the neural loci of intentions are dependent on the nature of the decision, which would make generalizing across paradigms difficult (Gilbert & Fung, 2018).

Indeed, our results can be seen as a warning when interpreting results from intention decoding. By using multivariate methods one can trace stable states in action decision processes. Our results show that the intuitive conclusion that these states are representations of action decisions is potentially problematic, and that the representational (i.e., stable) character of the process may only hold within a certain range of factors. In our experiment, the stable (and possibly representational) character of intentional processes disappeared when we varied the context in which the decision was made. Other, thus far unexplored factors could potentially have a similar effect. In more general terms, the invariance that is found during multivariate decoding may be dependent on invariance in the experimental setting.

To conclude, the processes leading to an action decision seem more context-dependent than it is often assumed. This may alter the way we think about the implementation of intentions, and the way they can be studied. More work is needed to assess whether intention are context-dependent through and through, or whether there is a context-invariant representational core that we have not discovered yet.

## Acknowledgements

SU was supported by a Marie Sklodowska-Curie grant (number 657605) from the European Union’s Framework Programme Horizon 2020. KG was funded by DFG grants GRK1589/1 and FK:JA945/3-1. JDH was funded by the Bernstein Computational Neuroscience Program of the German Federal Ministry of Education and Research BMBF (Grant 01GQ0411) and by the Excellence Initiative of the German Federal Ministry of Education and Research and DFG (Grants GSC86/1-2009, KFO247, HA 5336/1-1, SFB 940 and JA 945/3- 1/SL185/1-1.) The authors wish to thank Nora Swaboda for help with data collection, and Tim van Mourik for statistical advice.

## Footnotes

1 Goals (and therefore intentions) can be defined at many different levels (Uithol et al., 2012; Uithol, van Rooij, Bekkering, & Haselager, 2011). Consequently, actions that are purposeless on one level (say the movement in a Libet experiment) can be attributed a goal at a higher level (e.g., completing the experiment or complying with the experimenter)

## References

Andersen, R. A., & Cui, H. (2009). Intention, action planning, and decision making in parietal-frontal circuits. Neuron, 63(5), 568–583. http://doi.org/10.1016/j.neuron.2009.08.028

Ansuini, C., Cavallo, A., Bertone, C., & Becchio, C. (2014). Intentions in the Brain: The Unveiling of Mister Hyde. The Neuroscientist. http://doi.org/10.1177/1073858414533827

Bennur, S., & Gold, J. I. (2011). Distinct representations of a perceptual decision and the associated oculomotor plan in the monkey lateral intraparietal area. The Journal of Neuroscience.

Brass, M., & Haggard, P. (2008). The What, When, Whether Model of Intentional Action. The Neuroscientist, 14(4), 319–325. http://doi.org/10.1177/1073858408317417

Bratman, M. E. (1987). Intention, plans, and practical reason. Cambridge, MA: Harvard University Press.

Chang, C. C., & Lin, C. J. (2011). LIBSVM: a library for support vector machines. ACM Transactions on Intelligent Systems and Technology TIST, 2(3), 27.

Cisek, P. (2007). Cortical mechanisms of action selection: the affordance competition hypothesis. Philosophical Transactions of the Royal Society B-Biological Sciences, 362(1485), 1585–1599. http://doi.org/10.1146/annurev.neuro.20.1.25

Clark, A. (1997). Being there: Putting body, brain, and world together again. Cambridge, MA: MIT Press.

Cole, M. W., Ito, T., & Braver, T. S. (2016). The Behavioral Relevance of Task Information in Human Prefrontal Cortex. Cerebral Cortex (New York, N.Y. : 1991), 26(6), 2497–2505. http://doi.org/10.1093/cercor/bhv072

Desmurget, M., & Sirigu, A. (2012). Conscious motor intention emerges in the inferior parietal lobule. Current Opinion in Neurobiology.

Desmurget, M., Reilly, K. T., Richard, N., Szathmari, A., Mottolese, C., & Sirigu, A. (2009). Movement Intention After Parietal Cortex Stimulation in Humans. Science, 324(5928), 811–813. http://doi.org/10.1126/science.1169896

Fedorenko, E., Duncan, J., & Kanwisher, N. (2013). Broad domain generality in focal regions of frontal and parietal cortex. Proceedings of the National Academy of Sciences, 110(41), 16616–16621. http://doi.org/10.1073/pnas.1315235110

Flandin, G., & Friston, K. J. (2017). Analysis of family-wise error rates in statistical parametric mapping using random field theory. Human Brain Mapping, 4, 417. http://doi.org/10.1002/hbm.23839

Fried, I., Katz, A., McCarthy, G., Sass, K. J., Williamson, P., Spencer, S. S., & Spencer, D. D. (1991). Functional organization of human supplementary motor cortex studied by electrical stimulation. The Journal of Neuroscience, 11(11), 3656–3666.

Friston, K. J., Holmes, A. P., Worsley, K. J., Poline, J. P., Frith, C. D., & Frackowiak, R. S. J. (2004). Statistical parametric maps in functional imaging: A general linear approach. Human Brain Mapping, 2(4), 189–210. http://doi.org/10.1002/hbm.460020402

Gallivan, J. P., Johnsrude, I. S., & Randall Flanagan, J. (2015). Planning Ahead: Object-Directed Sequential Actions Decoded from Human Frontoparietal and Occipitotemporal Networks. Cerebral Cortex, 1–23. http://doi.org/10.1093/cercor/bhu302

Gallivan, J. P., McLean, D. A., Smith, F. W., & Culham, J. C. (2011a). Decoding Effector-Dependent and Effector-Independent Movement Intentions from Human Parieto-Frontal Brain Activity. The Journal of Neuroscience, 31(47), 17149–17168. http://doi.org/10.1523/JNEUROSCI.1058-11.2011

Gallivan, J. P., McLean, D. A., Valyear, K. F., Pettypiece, C. E., & Culham, J. C. (2011b). Decoding Action Intentions from Preparatory Brain Activity in Human Parieto-Frontal Networks. The Journal of Neuroscience, 31(26), 9599–9610. http://doi.org/10.1523/JNEUROSCI.0080-11.2011

Gibson, J. (1979). The ecological approach to visual perception. Boston: Houghton Mifflin.

Gilbert, S. J., & Fung, H. (2018). Decoding intentions of self and others from fMRI activity patterns. NeuroImage, 172, 278–290. http://doi.org/10.1016/j.neuroimage.2017.12.090

Goldberg, G. (1985). Supplementary Motor Area Structure and Function - Review and Hypotheses. Behavioral and Brain Sciences, 8(4), 567–588. http://doi.org/10.1017/S0140525X00045167

Görgen, K., Hebart, M. N., Allefeld, C., & Haynes, J.-D. (2017). The same analysis approach: Practical protection against the pitfalls of novel neuroimaging analysis methods. NeuroImage. http://doi.org/10.1016/j.neuroimage.2017.12.083

Grafton, S. T., & Hamilton, A. F. de C. (2007). Evidence for a distributed hierarchy of action representation in the brain. Human Movement Science, 26(4), 590–616.

Haggard, P. (2005). Conscious intention and motor cognition. Trends in Cognitive Sciences, 9(6), 290–295.

Haggard, P. (2008). Human volition: towards a neuroscience of will. Nature Reviews: Neuroscience, 9(12), 934–946. http://doi.org/10.1038/nrn2497

Hamilton, A. F. de C., & Grafton, S. T. (2007). The motor hierarchy: from kinematics to goals and intentions. In P. Haggard, Y. Rossetti, & M. Kawato (Eds.), Attention & Performance 22. Sensorimotor Foundations of Higher Cognition Attention and Performance (pp. 381–408). Oxford: Oxford University Press.

Haynes, J.-D., Sakai, K., Rees, G., Frith, C. D., & Passingham, R. E. (2007). Reading Hidden Intentions in the Human Brain. Current Biology, 17(4), 323–328. http://doi.org/10.1016/j.cub.2006.11.072

Hebart, M. N., Görgen, K., & Haynes, J.-D. (2015). The Decoding Toolbox (TDT): a versatile software package for multivariate analyses of functional imaging data. Frontiers in Neuroinformatics, 8(174). http://doi.org/10.3389/fninf.2014.00088

Jahanshahi, M. (1998). Willed Actions and it Impairments. Cognitive Neuropsychology, 15(6-8), 483–533. http://doi.org/10.1080/026432998381005

James, W. (1890). The principles of psychology. New York: H. Holt and company.

JASP Team. (2017). JASP (version 0.8.1.1).

Kornhuber, H. H., & Deecke, L. (1965). Hirnpotentialänderungen bei Willkürbewegungen und passiven Bewegungen des Menschen: Bereitschaftspotential und reafferente Potentiale. Pflügers Archiv Für Die Gesamte Physiologie Des Menschen Und Der Tiere, 284(1), 1–17. http://doi.org/10.1007/BF00412364

Kriegeskorte, N., Simmons, W. K., Bellgowan, P. S. F., & Baker, C. I. (2009). Circular analysis in systems neuroscience: the dangers of double dipping. Nature Neuroscience, 12(5), 535–540. http://doi.org/10.1038/nn.2303

Krieghoff, V., Brass, M., Prinz, W., & Waszak, F. (2009). Dissociating what and when of intentional actions. Frontiers in Human Neuroscience, 3(3), 1–10. http://doi.org/10.3389/neuro.09.003.2009

Lee, M. D., & Wagenmakers, E.-J. (2014). Bayesian Cognitive Modeling.

Momennejad, I., & Haynes, J.-D. (2012). Human anterior prefrontal cortex encodes the ‘what’and “when”of future intentions. NeuroImage, 61(1), 139–148. http://doi.org/10.1016/j.neuroimage.2012.02.079

Mylopoulos, M., & Pacherie, E. (2017). Intentions and Motor Representations: the Interface Challenge. Review of Philosophy and Psychology, 8(2), 317–336. http://doi.org/10.1007/s13164-016-0311-6

Oldfield, R. C. (1971). The assessment and analysis of handedness: The Edinburgh inventory. Neuropsychologia, 9(1), 97–113. http://doi.org/10.1016/0028-3932(71)90067-4

Pacherie, E. (2008). The phenomenology of action: A conceptual framework. Cognition, 107(1), 179–217.

Peirce, J. W. (2007). PsychoPy—Psychophysics software in Python. Journal of Neuroscience Methods, 162(1-2), 8–13. http://doi.org/10.1016/j.jneumeth.2006.11.017

Schurger, A. A., & Uithol, S. (2015). Nowhere and Everywhere: The Causal Origin of Voluntary Action. Review of Philosophy and Psychology, 6(4), 761–778. http://doi.org/10.1007/s13164-014-0223-2

Sigala, N., Kusunoki, M., Nimmo-Smith, I., Gaffan, D., & Duncan, J. (2008). Hierarchical coding for sequential task events in the monkey prefrontal cortex. Proceedings of the National Academy of Sciences, 105(33), 11969–11974.

Soon, C. S., Brass, M., Heinze, H.-J., & Haynes, J.-D. (2008). Unconscious determinants of free decisions in the human brain. Nature Neuroscience, 11(5), 543–545. http://doi.org/10.1038/nn.2112

Uithol, S., Burnston, D., & Haselager, W. F. G. (2014). Why we may not find intentions in the brain. Neuropsychologia, 56, 129–139. http://doi.org/10.1016/j.neuropsychologia.2014.01.010

Uithol, S., van Rooij, I., Bekkering, H., & Haselager, W. F. G. (2011). Understanding motor resonance. Social Neuroscience, 6(4), 388–397. http://doi.org/10.1080/17470919.2011.559129

Uithol, S., van Rooij, I., Bekkering, H., & Haselager, W. F. G. (2012). Hierarchies in Action and Motor Control. Journal of Cognitive Neuroscience, 24(5), 1077–1086. http://doi.org/10.1162/jocn_a_00204

Vul, E., Harris, C., Winkielman, P., & Pashler, H. (2009). Puzzlingly High Correlations in fMRI Studies of Emotion, Personality, and Social Cognition. Perspectives on Psychological Science, 4(3), 274–290. http://doi.org/10.1111/j.1745-6924.2009.01125.x

Wilson, M. (2002). Six views of embodied cognition. Psychonomic Bulletin & Review, 9(4), 625–636. http://doi.org/10.3758/BF03196322

Wokke, M. E., Knot, S. L., Fouad, A., & Ridderinkhof, K. R. (2016). Conflict in the kitchen: Contextual modulation of responsiveness to affordances. Consciousness and Cognition, 40(C), 141–146. http://doi.org/10.1016/j.concog.2016.01.007

Woolgar, A., Thompson, R., Bor, D., & Duncan, J. (2011). Multi-voxel coding of stimuli, rules, and responses in human frontoparietal cortex. NeuroImage, 56(2), 744–752. http://doi.org/10.1016/j.neuroimage.2010.04.035

Zapparoli, L., Seghezzi, S., Scifo, P., Zerbi, A., Banfi, G., Tettamanti, M., & Paulesu, E. (2018). Dissecting the neurofunctional bases of intentional action. Proceedings of the National Academy of Sciences, 11, 201718891. http://doi.org/10.1073/pnas.1718891115

